# Differential effects of inducible cerebellar granule cell and Purkinje cell ablation on motor coordination and motor learning in adult mice

**DOI:** 10.1101/2025.09.05.674420

**Authors:** Eiko Kato, Misato Yasumura, Shigeyoshi Itohara, Kenji Sakimura, Takeshi Uemura, Masayoshi Mishina

## Abstract

The cerebellum regulates motor coordination and motor learning through highly organized circuits composed mainly of granule cells (GCs) and Purkinje cells (PCs). To investigate their distinct roles, we generated two lines of inducible transgenic mice in which either GCs or PCs could be selectively ablated in adulthood by administration of the progesterone receptor antagonist RU-486. This system combined a Cre recombinase-progesterone receptor fusion, in which Cre activity is induced in an RU-486-dependent manner, with a Cre-dependent diphtheria toxin A expression to achieve cell-type-specific ablation. High-dose RU-486 induced nearly complete loss of either GCs or PCs and resulted in severe ataxia. When partial ablation was induced by low-dose RU-486, different phenotypes emerged. Mice retaining about 20% of PCs were still able to improve motor coordination in the rotarod test and maintained performance in the balance beam test comparable to that of controls. In contrast, mice with about 30% of GCs remaining showed marked deficits, failing to improve across rotarod trials and exhibiting reduced latency to fall in the balance beam test. These results suggest that while both GCs and PCs are indispensable for motor coordination, a sufficient number of GCs is required for both motor coordination and motor learning. This inducible ablation model highlights the differential contributions of cerebellar neurons and provides a valuable tool to dissect circuit-specific functions in the adult brain.

## Introduction

The cerebellum is essential for the precise regulation of motor coordination and motor learning (Boyden et al., 2004; Christian et al., 2003). The cerebellar cortex is highly organized, consisting mainly of granule cells (GCs) and Purkinje cells (PCs). Mossy fibers from the cell bodies in the spinal cord and brainstem provide excitatory input to GCs, whose axons form parallel fibers that synapse onto the distal dendrites of PCs. In contrast, climbing fibers from the inferior olive innervate the proximal dendrites of PCs. PCs function as the sole output neurons of the cerebellar cortex (Altman & Bayer, 1997). These features make the cerebellum an ideal model for investigating brain wiring and neuronal function.

Investigation of the roles of specific neurons or neuronal pathways is essential for a comprehensive understanding of brain function. Classical mouse mutants with cerebellar dysfunction, such as lurcher, hot-foot, staggerer, Purkinje cell degeneration (pcd), and weaver, exhibit cerebellar atrophy, ataxia, and deficits in motor coordination (Lalonde & Strazielle, 2007). Both lurcher and pcd mutants suffer from almost complete loss of PCs in addition to severe GC death (Caddy & Biscoe, 1979; Heckroth & Eisenman, 1991; Mullen et al., 1976; Landis & Mullen, 1978). In contrast, staggerer and weaver mutants show partial PC loss of approximately 75% and 45%, respectively (Herrup, 1983; Herrup & Mullen, 1979; Blatt & Eisenman, 1985a,b). Although the motor coordination of these mutants has been extensively studied, their developmental abnormalities and combined neuronal loss complicate interpretation of cell-type–specific functions.

To overcome these limitations, cell-type-specific and inducible ablation of neurons in the adult brain provides a powerful approach for in vivo functional analysis. Here, we generated two lines of transgenic mice in which selective ablation of either GCs or PCs can be induced in adulthood by oral administration of the progesterone receptor antagonist RU-486. Using these models, we demonstrate that partial ablation of GCs and PCs differentially affects motor coordination and motor learning, thereby revealing distinct contributions of cerebellar neuron types in adult brain function.

## Materials and Methods

### Animals

Transgenic mice in which cerebellar GCs can be selectively ablated by drug administration were generated by crossing Eno2-STOP-DTA homozygous mice (Kobayakawa et al., 2007; Kishioka et al., 2009) with heterozygous *GluN2c-CrePR* mice (ECP25 line, Tsujita et al., 1999). Transgenic mice in which cerebellar PCs can be selectively ablated were generated by crossing Eno2-STOP-DTA homozygous mice with heterozygous *GluD2-CreP*R mice expressing a Cre recombinase–progesterone receptor fusion protein (CrePR) under the control of *GluD2* promoter (D2CPR mice line, Kitayama et al., 2001). All animal procedures were approved by the Animal Care and Use Committee of the Graduate School of Medicine, the University of Tokyo (Approval #1721T062), and all experiments were conducted in accordance with the Guidelines for the Care and Use of Laboratory Animals of the University of Tokyo.

### Drug treatment

RU-486 (Sigma) was dissolved in milk containing 20% (v/v) ethanol to yield final concentrations of 15, 3, or 0.5 mg/ml, and was administered orally to mice at doses of 0.3, 0.06, or 0.01 mg/g body weight at postnatal week 7.

### Immunohistochemistry and image acquisition

Under deep pentobarbital anesthesia, mice were perfused transcardially with 4% paraformaldehyde in 0.1 M sodium phosphate buffer (pH 7.2). Immnohistochemical analysis was perfeomed as described previously (Uemura et al., 2010). Briefly, parasagittal cerebellar sections (50 μm thick) were prepared using a VT1000S microslicer (Leica Microsystems). After blocking with 10% donkey serum, the sections were incubated with rabbit anti-calbindin (Nakagawa et al., 1998) and mouse anti-neuronal nuclei (NeuN) antibodies (Chemicon International), followed by incubating with Alexa 488 conjugated anti-rabbit and Alexa 555 conjugated anti-mouse IgG antibodies (Invitrogen). Images were taken with an AX-70 microscope equipped with a DP11 CCD camera (Olympus). The numbers of NeuN-positive GCs were counted per 0.01 mm² in the GC layer of lobules IV/V and X. The numbers of calbindin-positive PCs were counted along the entire PC layer in midsagittal cerebellar sections. Quantification was performed using NIH ImageJ software.

### Behavioral analysis

Footprints were obtained 7 days after RU-486 administration by dipping the hind feet in Indian ink and allowing the mice to walk across white paper. The rotating rod tests were performed with an apparatus consisting of a 3.2 cm-diameter rod (RRSW-3002, O’Hara). The constant-speed rotarod test was first conducted 7 days after RU-486 administration at various speeds. The rod speed was set from 1 to 40 rpm in 3-rpm increments, with intertrial intervals of 30 min. A cutoff time of 2 min was applied for all tests, and the latency to fall was recorded. This was defined as the 1st trial. The same procedure was repeated on days 14 and 21 after RU-486 administration, corresponding to the 2nd and 3rd trials, respectively. The 1st, 2nd, and 3rd trials were performed sequentially using the same cohort of mice. The balance beam test was performed as previously described (Airaksinen et al., 1997) using a beam 15 mm in diameter, 50 cm in length, and elevated 40 cm above the floor (TR-3002; O’Hara & Co.). Mice were placed at the midpoint of the rod, and the latency to fall was recorded. The cutoff time was set at 60 s. Each mouse received six trials per day with 20-min intertrial intervals for 2 days before and 14 days after RU-486 administration.

### Statistical analysis

Statistical significance was evaluated using two-way repeated measures ANOVA. A *p* value < 0.05 was considered statistically significant.

## Results

### Generation of inducible mouse lines for cerebellar GC or PC ablation

To investigate the roles of cerebellar GCs and PCs in adult motor coordination and motor learning, we generated two transgenic mouse lines in which either GCs or PCs can be selectively ablated by drug administration, through a combination of an inducible Cre-mediated recombination system and Cre-dependent expression of the *diphtheria toxin A* (*DTA*) gene. For cerebellar GC-specific ablation, *Eno2-STOP-DTA* mice were crossed with GC-specific CrePR-expressing *GluN2c-CrePR* mice, generating *GluN2c -CrePR*; *Eno2-STOP-DTA* mice, hereafter referred to as GC-specific DTA (Fig. 1A). For PC-specific ablation, *Eno2-STOP-DTA* mice were crossed with PC-specific CrePR-expressing *GluD2*-CrePR mice, generating *GluD2-CrePR*; *Eno2-STOP-DTA* mice, hereafter referred to as PC-specific DTA (Fig. 1B).

**Fig. 1.**
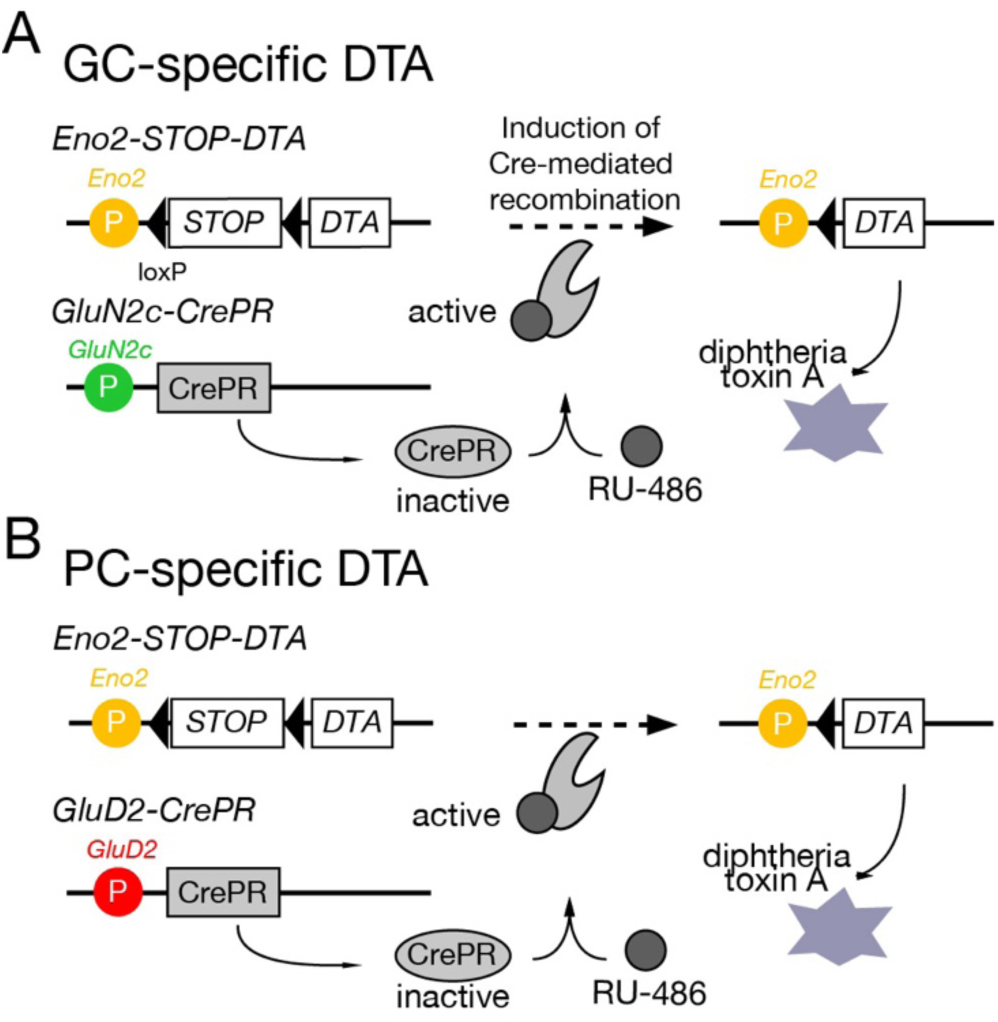
Inducible and specific ablation of cerebellar neurons by CrePR-loxP recombination system. (A and B) Inducible ablation of cerebellar GCs (A) and PCs (B) by antiprogestin RU-486 administration. *Eno2-STOP-DTA* mice carry a Cre-inducible DTA gene in the *Eno2* locus. *GluN2c-CrePR* and *GluD2-CrePR* lines express CrePR under *GluN2c* or *GluD2* promoter, respectively. RU-486 activates CrePR, excising the floxed stop codon and inducing DTA expression, resulting in GC- or PC-specific ablation in the adult cerebellum. GC, granule cell; PC, Purkinje cell; DTA, diphtheria toxin A.

### High-efficiency cerebellar GC ablation by RU-486 administration in GC-specific DTA mice

To induce CrePR recombinase activity, RU-486 was administered orally at a dose of 0.3 mg/g body weight to GC-specific DTA mice at postnatal week 7, and the same administration schedule was applied throughout all experiments. *Eno2-STOP-DTA* mice treated with RU-486 served as control, and the same control group was used in all subsequent experiments. To evaluate the extent of GC ablation, immunohistochemical analyses using anti-calbindin and NeuN antibodies were performed at 3, 5, and 7 days after RU-486 administration (Fig. 2A). In RU-486–treated GC-specific DTA mice, the number of NeuN-positive GCs gradually decreased from day 3 to day 7, reaching approximately 13% of that in control mice in lobules IV/V at day 7 (Fig. 2B). In contrast, no reduction of GCs was observed in lobule X (Fig. 2C). This observation was consistent with previous reports showing that RU-486–induced Cre activity in *GluN2c* -CrePR mice is absent in lobule X (Tujita et al., 1999). Thus, oral administration of RU-486 at 0.3 mg/g body weight resulted in the ablation of nearly 90% of cerebellar GCs in GC-specific DTA mice. These mice are hereafter referred to as GC-high ablation mice.

**Fig. 2.**
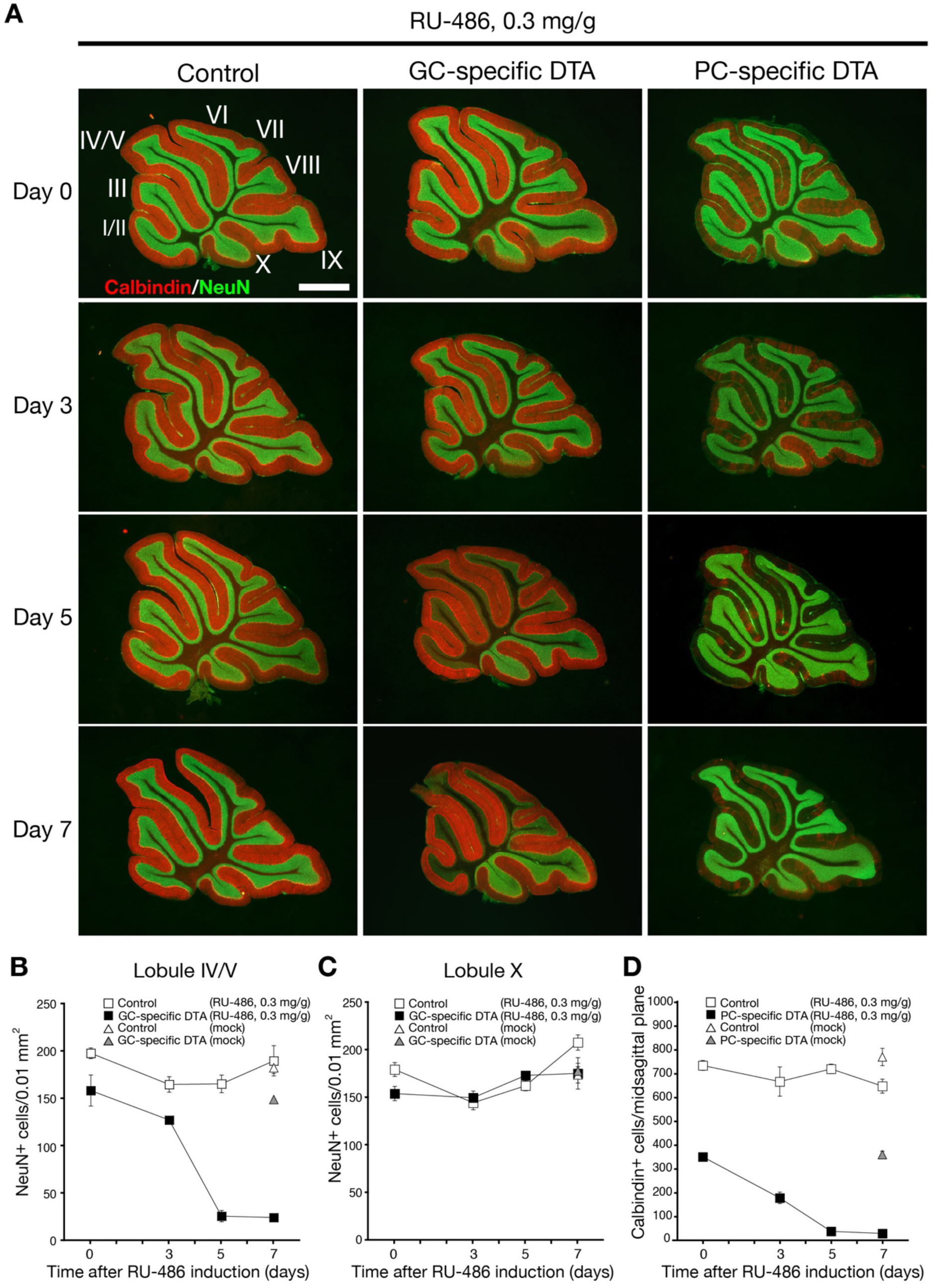
Extensive ablation of cerebellar GCs or PCs induced by high-dose RU-486 administration. (A) Immunohistochemical images of NeuN (green) and calbindin (red) in control, GC-DTA, and PC-DTA mice before and at 3, 5, and 7 days after RU-486 administration at 0.3 mg/g body weight at 7 weeks of age. (B, C) Changes in NeuN^+^ GC density in lobules IV/V (B) and X (C) of GC-DTA and control mice after RU-486 administration. (D) Changes in calbindin^+^ PC density of PC-DTA and control mice after RU-486 administration. PCs were quantified in midsagittal sections. All values presented as mean ± SEM. n = 3 mice per group. Scale bar, 1 mm. I–X indicate cerebellar folia I–X.

### High-efficiency cerebellar PC ablation by RU-486 administration in PC-specific DTA mice

To ablate PCs in the adult mice, RU-486 was administered orally at a dose of 0.3 mg/g body weight to PC-specific DTA mice. Cerebellar sections obtained after RU-486 administration were stained with anti-calbindin and NeuN antibodies (Fig. 2A). In PC-specific DTA mice, the number of calbindin-positive PCs was already reduced to approximately 48% of control levels before RU-486 administration. After RU-486 treatment, the number of PCs progressively decreased from day 3 to day 5 and reached approximately 5% of control mice by day 7 (Fig. 2D). Thus, oral administration of RU-486 at 0.3 mg/g body weight resulted in the ablation of nearly 95% of PCs in PC-specific DTA mice. These mice are hereafter referred to as PC-high ablation mice.

### GC- and PC-high ablation mice exhibited severe ataxic gait with slight differences

GC-high ablation and PC-high ablation mice exhibited severe ataxic gait. Both groups used the entire hind paws to walk and maintain balance, whereas control mice tended to use only the toes and forepart of the foot. However, subtle differences in motor phenotypes were observed. GC-high ablation mice sometimes scrambled on their hind limbs, which was not observed in PC-high ablation mice. Differences were also evident in the footprint pattern (Fig. 3A). We next assessed motor performance using the constant-speed rotarod test. Both GC- and PC-high ablation mice exhibited severe impairments in motor coordination as assessed by the constant-speed rotarod test. Latency to fall in the 1st, 2nd and 3rd trials after RU-486 administration were significantly reduced in GC-high ablation mice compared with control mice (1st, p = 0.0001; 2nd, p < 0.0001; 3rd, p < 0.0001; ANOVA with repeated measures, group × session) (Fig. 3B and 3C). Similarly, latency to fall in PC-high ablation mice were significantly reduced compared with control mice (1st, p = 0.0102; 2nd, p = 0.0105; 3rd, p = 0.0022; ANOVA with repeated measures, group × session) (Fig. 3B and 3D). In contrast, no significant differences were observed between GC-high ablation and PC-high ablation mice in the 1st trial (Fig. 3C and 3D). Interestingly, in the 2nd and 3rd trial, PC-high ablation mice exhibited higher latency to fall than GC-high ablation mice (p = 0.0488 and p < 0.0001, respectively; ANOVA with repeated measures, group × session).

**Fig. 3.**
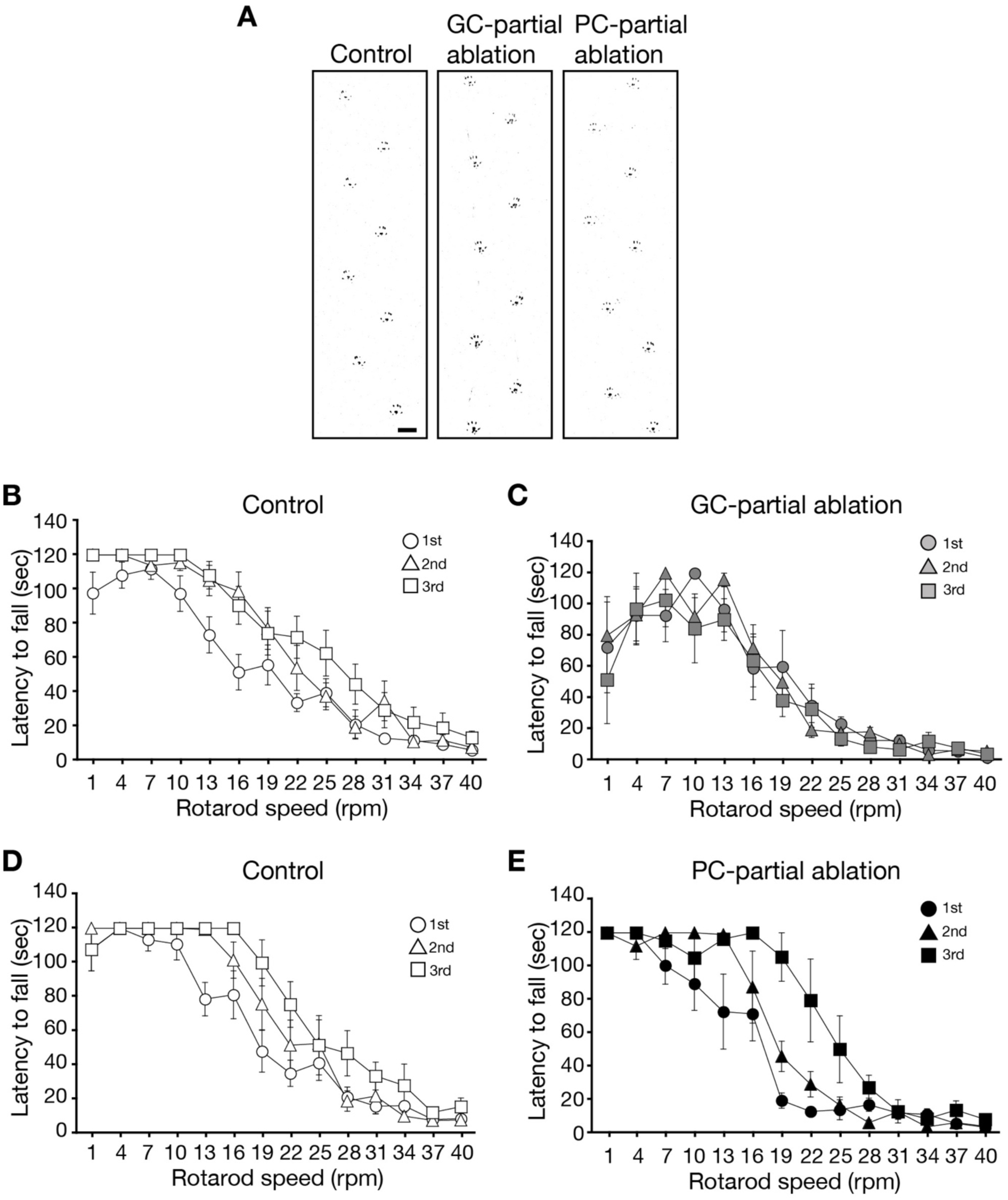
Motor coordination in GC- and PC-high ablation mice in rotarod test. (A) Representative footprint patterns of control (left), GC-high ablation (middle), and PC-high ablation (right) mice 7 days after RU-486 administration (0.3 mg/g). (B–D) Rotarod test. Mice underwent the first trial on day 7, the second trial on day 14, and the third trial on day 21 after RU-486 administration (0.3 mg/g body weight). All values presented as mean ± SEM. Control in (B), n = 17; GC-high ablation in (C), n = 7; PC-high ablation in (D), n = 4.

### Partial ablation of cerebellar GC ablation by RU-486 administration in GC-specific DTA mice

To examine how the extent of GC ablation affects motor coordination, we orally administered RU-486 at a low dose of 0.06 mg/g body weight to GC-specific DTA mice. To evaluate the extent of GC ablation, immunohistochemical analyses using anti-calbindin and NeuN antibodies were performed after RU-486 administration (Fig 4A). In RU-486–treated GC-specific DTA mice, the number of NeuN-positive GCs in lobules IV/V decreased by day 5 and then showed little change through day 21, resulting in approximately 30% of control mice (Fig. 4B). In contrast, no reduction of GCs was observed in lobule X (Fig. 4C). Thus, oral administration of RU-486 at 0.06 mg/g body weight induced partial ablation of GCs, eliminating nearly 70% of cells in lobules IV/V. These mice are hereafter referred to as GC-partial ablation mice.

**Fig. 4.**
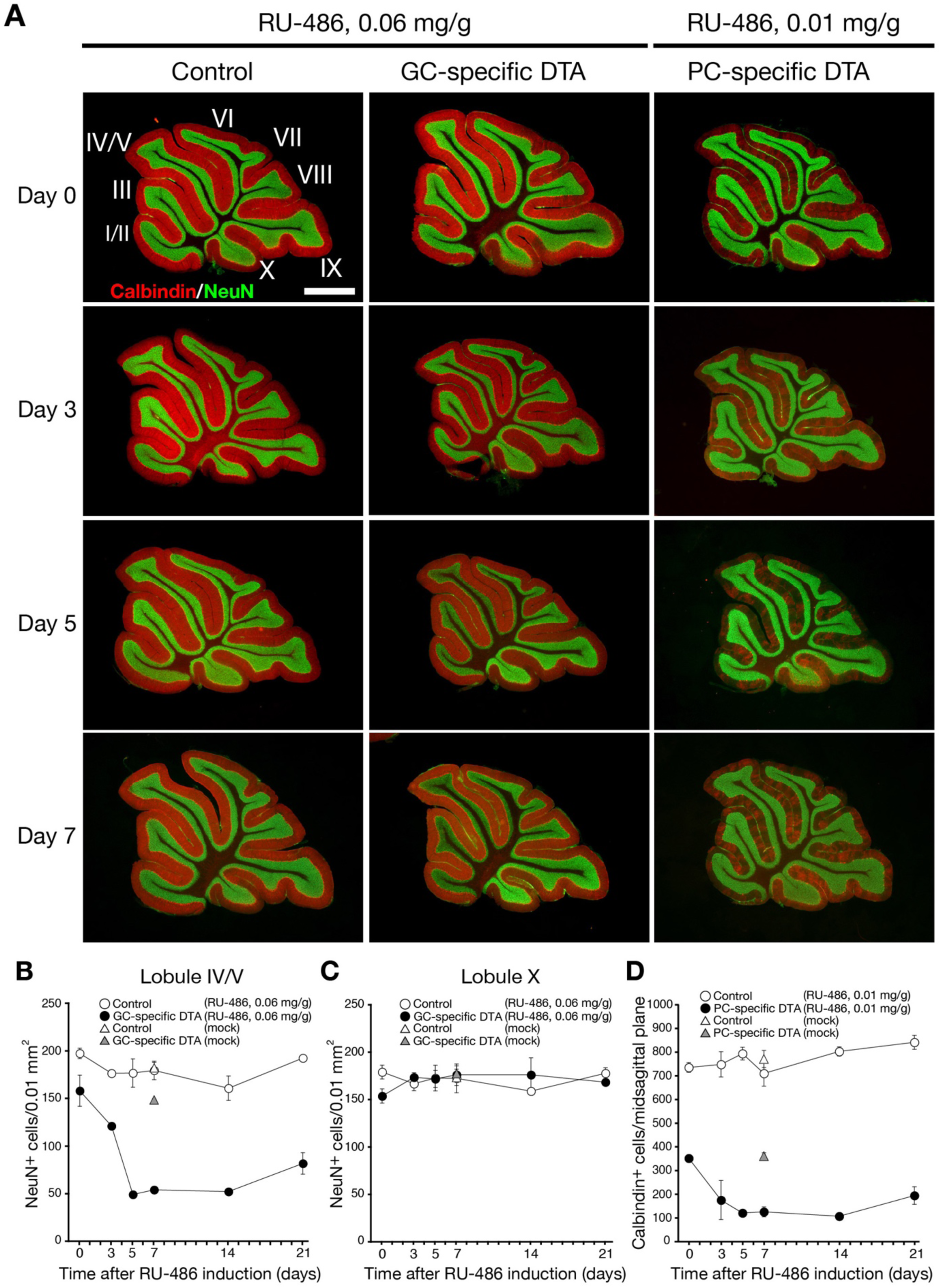
Partial ablation of cerebellar GCs or PCs induced by low-dose RU-486 administration. (A) Immunohistochemical images of NeuN (green) and calbindin (red) in control, GC-partial ablation, and PC-partial ablation mice before and at 3, 5, and 7 days after RU-486 administration (0.06 mg/g body weight for GC-partial ablation and 0.01 mg/g body weight for PC-partial ablation) at 7 weeks of age. (B, C) Changes in NeuN^+^ GC density in lobules IV/V (B) and X (C) of GC-partial ablation and control mice after RU-486 administration. (D) Changes in calbindin^+^ PC density of PC-partial ablation and control mice after RU-486 administration. PCs were quantified in midsagittal sections. All values presented as mean ± SEM. n = 3 mice per group. Scale bar, 1 mm. I–X indicate cerebellar folia I–X.

### Partial ablation of cerebellar PC ablation by RU-486 administration in PC-specific DTA mice

To examine how the extent of PC ablation affects motor coordination, PC-specific DTA mice were orally administrated RU-486 at low dose of 0.01 mg/g body weight. Cerebellar sections were immunostained with anti-calbindin and NeuN antibodies (Fig. 4A). In RU-486–treated PC-specific DTA mice, the number of calbindin-positive PCs decreased by day 5 and then showed little change through day 21, resulting in approximately 20% of control mice (Fig. 4D). Thus, oral administration of RU-486 at 0.01 mg/g body weight resulted in the ablation of nearly 80% of PCs in PC-specific DTA mice. These mice are hereafter referred to as PC-partial ablation mice.

### GC- and PC-partial ablation differentially impair motor coordination and motor learning

Both GC-partial ablation mice and PC-partial ablation mice exhibited ataxic gait, similar to GC-high ablation and PC-high ablation mice. However, the severity of ataxic gait was milder in the partially ablated mice. Although both GC-partial and PC-partial ablation mice showed ataxic gait, they could walk in a straight line, and their footprint patterns were indistinguishable from those of control mice (Fig. 5A).

**Fig. 5.**
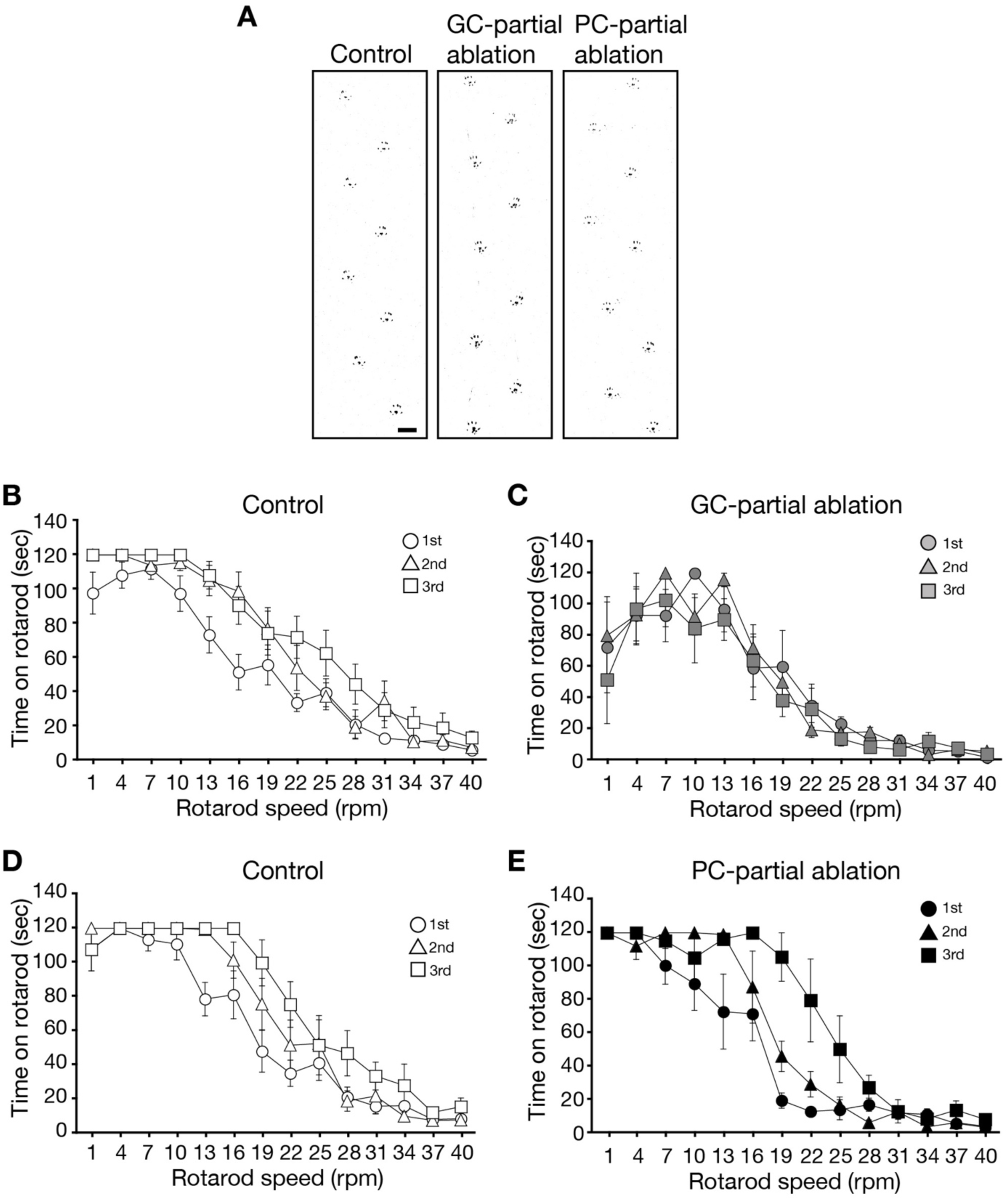
Motor coordination in GC- and PC-partial ablation mice in rotarod test. (A) Representative footprint patterns of control (left), GC-partial ablation (middle), and PC-partial ablation (right) mice 7 days after RU-486 administration (0.06 mg/g body weight for control and GC-partial ablation, and 0.01 mg/g for PC-partial ablation). (B–E) Rotarod test. Mice underwent the first trial on day 7, the second trial on day 14, and the third trial on day 21 after RU-486 administration. RU-486 was given at 0.06 mg/g body weight in control (B) and GC-partial ablation (C) mice, and at 0.01 mg/g body weight in control (D) and PC-partial ablation (E) mice. All values presented as mean ± SEM. Control in (B), n = 13; GC-DTA in (C), n = 7; control in (D), n = 9; PC-DTA in (D), n = 4.

To further examine the motor coordination of these mutant mice, we performed the constant-speed rotarod test as described in Fig. 3. In the 1st trial, GC-partial ablation mice and PC-partial ablation mice exhibited latency to fall comparable to those of their respective control mice (p = 0.5030 and p = 0.5984, respectively; ANOVA with repeated measure, group × session) (Fig. 5B-D). Interestingly, control mice showed a significant increase in latency to fall from the 1st to the 3rd trial (p < 0.0001; ANOVA with repeated measures, trial effect) (Fig. 5B). In contrast, GC-partial ablation mice did not exhibit such an increase in latency to fall (Fig. 5C) (p = 0.3096; ANOVA with repeated measures, trial effect). Conversely, PC-partially ablated mice exhibited an increase in latency to fall from the 1st to the 3rd trial, similar to that observed in control mice (Fig. 5D and 5E) (p < 0.0001 for control; p < 0.0001 for PC-partial ablation mice; ANOVA with repeated measures, trial effect). However, their latency to fall was reduced in the first trial (p = 0.0108; two-way ANOVA, group × session). These findings suggest that while PC-partial ablation mice retain the ability for motor learning in the variable-speed rotarod test, such adaptive learning is impaired in GC-partial ablation mice.

To further analyze the differences in motor coordination between GC-partial ablation and PC-partial ablation mice, we conducted the balance beam test (Fig. 6). Control mice consistently remained on the rod across trials from day 0 (the day of RU-486 administration) to day 14. In contrast, GC-partial ablation mice exhibited a sharp reduction in latency to fall on day 5, followed by a gradual increase up to day 14 (Fig. 6A). The abrupt decline in latency to fall observed on day 5 corresponded closely to the time course of GC ablation (Fig. 4A). Although latency to fall gradually improved thereafter, they remained significantly lower than those of control mice (p < 0.001; ANOVA with repeated measure, group × session). Conversely, PC-partial ablation mice displayed latency to fall comparable to those of control mice throughout the testing period (p =0.2426; ANOVA with repeated measures, group × session) (Fig. 6B). These results suggested that partial ablation of GCs and PCs differentially affects motor coordination.

**Fig. 6.**
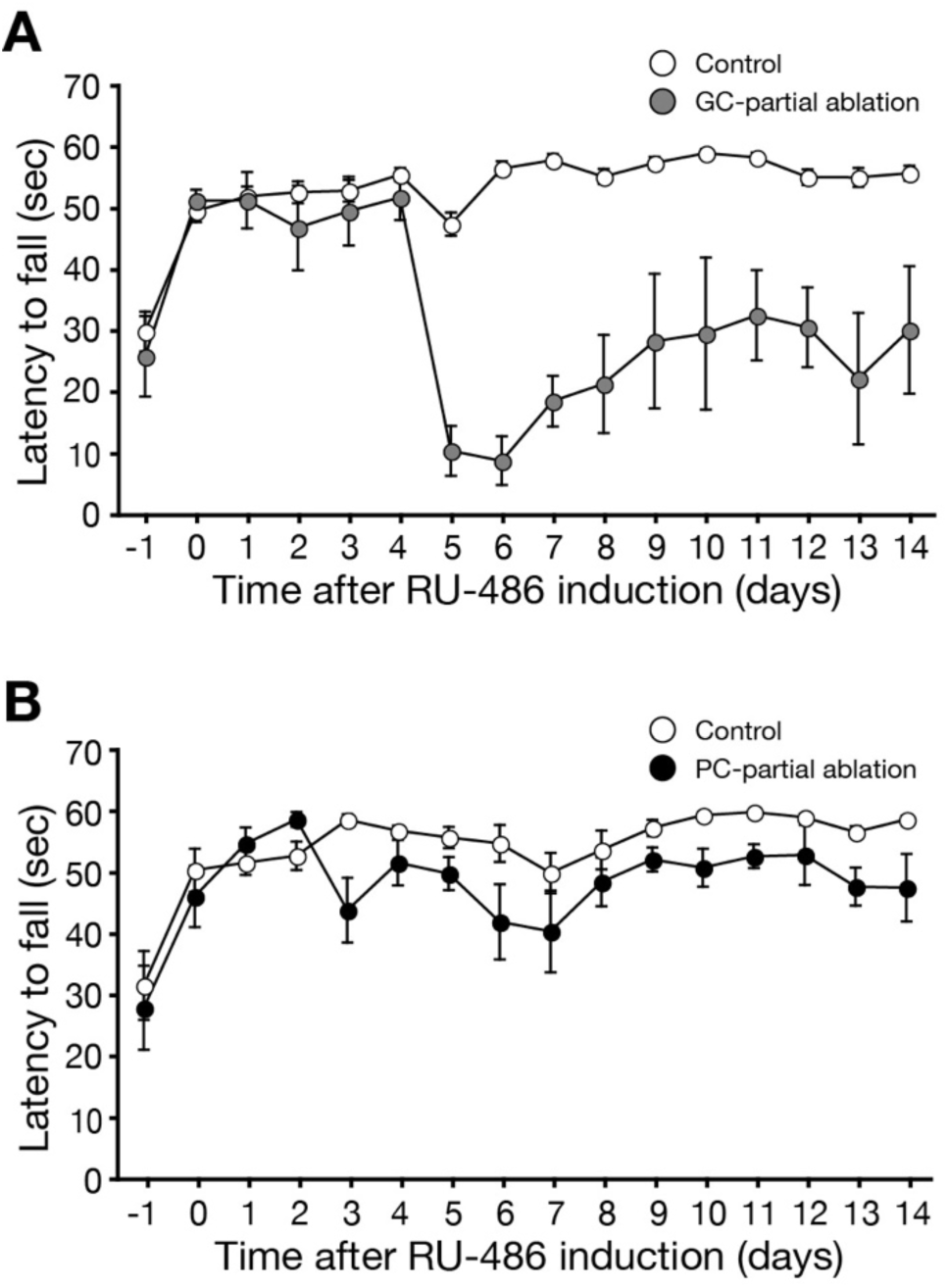
Motor coordination in GC- and PC-partial ablation mice in balance beam test. (A) Latency to fall from the beam in GC-partial ablation and control mice. RU-486 was administered at 0.06 mg/g body weight. Measurements were performed from 1 day before to 14 days after administration. Control, n = 16; GC-partial ablation, n = 5. (B) Latency to fall from the beam in PC-partial ablation and control mice. RU-486 was administered at 0.01 mg/g body weight. Measurements were performed from 1 day before to 14 days after administration. Control, n = 11; PC-partial ablation, n = 5. All values presented as mean ± SEM.

## Discussion

In this study, we established an inducible and selective ablation system that eliminates cerebellar GCs or PCs in adult mice by oral administration of RU-486. High-dose treatment caused near-complete loss of either GCs or PCs, resulting in severe ataxia, consistent with their essential roles in motor coordination (Palay & Chan-Palay, 1974). Importantly, low-dose RU-486 induced partial ablation, enabling evaluation of the distinct requirements of each cell type for motor coordination and motor learning.

Under partial ablation, clear differences emerged between GC- and PC-mutant mice. Mice retaining approximately 20% of PCs improved in the variable-speed rotarod test and performed normally in the balance beam test (Figs. 5 and 6), suggesting preserved motor learning. In contrast, mice with approximately 30% of GCs remaining failed to improve on the rotarod (Fig. 5) and showed reduced latency to fall in the beam test (Fig. 6). These findings suggest that motor learning requires a sufficient number of GCs, whereas a relatively small population of PCs can sustain adaptive motor improvement.

This difference likely reflects the organization of cerebellar circuitry. PCs integrate two excitatory inputs: parallel fibers from numerous GCs and a single climbing fiber from the inferior olive. Loss of GCs strongly disturbs the balance between parallel fiber and climbing fiber inputs, impairing motor learning (Napper & Harvey, 1988; Strata & Rossi, 1998). In contrast, reduced PC numbers may preserve both inputs in the remaining cells, maintaining conditions for learning despite diminished output.

Spontaneous mutants such as lurcher, pcd, staggerer, and weaver have provided important insights into cerebellar function but are complicated by developmental abnormalities (Caddy & Biscoe, 1979; Heckroth & Eisenman, 1991; Mullen et al., 1976; Landis & Mullen, 1978; Herrup, 1983; Herrup & Mullen, 1979; Blatt & Eisenman, 1985a,b; Lalonde & Strazielle, 2007). However, developmental abnormalities make interpretation difficult. Our inducible system complements these studies by enabling selective ablation of GCs or PCs in adults without developmental confounds.

In conclusion, both GCs and PCs are indispensable for motor coordination, but motor learning critically depends on maintaining a sufficient number of GCs. The inducible ablation system described here provides a valuable model for dissecting cell-type–specific contributions to cerebellar function in the adult brain.

## Acknowledgements

This work was supported in part by research grants from the Ministry of Education, Culture, Sports, Science, and Technology of Japan. We thank M. Watanabe for antibody against calbindin.

## Authors contributions

E.K. performed immunohistochemical analyses and behavioral experiments. M.Y. supervised the behavioral experiments. S.I. provided *Eno2-STOP-DTA* mice, and K.S. provided *GluN2c-CrePR* and *GluD2-CrePR* mice. E.K. and T.U. designed the study and wrote the manuscript. M.Y. and M.M. contributed to manuscript editing, and M.M. supervised the overall project.

